# Genetic modification of the shikimate pathway to reduce lignin content in switchgrass (*Panicum virgatum* L.) significantly impacts plant microbiomes

**DOI:** 10.1101/2024.05.02.592240

**Authors:** Shuang Liu, Ming-Yi Chou, Gian Maria Niccolò Benucci, Aymerick Eudes, Gregory Bonito

## Abstract

Switchgrass (*Panicum virgatum* L.) is considered a sustainable biofuel feedstock, given its fast-growth, low input requirements, and high biomass yields. Improvements in bioenergy conversion efficiency of switchgrass could be made by reducing its lignin content. Engineered switchgrass that expresses a bacterial 3-dehydroshikimate dehydratase (QsuB) has reduced lignin content and improved biomass saccharification due to the rerouting of the shikimate pathway towards the simple aromatic protocatechuate at the expense of lignin biosynthesis. However, the impacts of this QsuB trait on switchgrass microbiome structure and function remains unclear. To address this, wildtype and QsuB engineered switchgrass were grown in switchgrass field soils and samples were collected from inflorescences, leaves, roots, rhizospheres, and bulk soils for microbiome analysis. We investigated how QsuB expression influenced switchgrass-associated fungal and bacterial communities using high-throughput Illumina MiSeq amplicon sequencing of ITS and 16S rDNA. Compared to wildtype, QsuB engineered switchgrass hosted different microbial communities in roots, rhizosphere, and leaves. Specifically, QsuB engineered plants had a lower abundance of arbuscular mycorrhizal fungi (AMF). Additionally, QsuB engineered plants had fewer *Actinobacteriota* in root and rhizosphere samples. These findings may indicate that changes in the plant metabolism impact both organismal groups similarly, or potential interactions between AMF and the bacterial community. This study enhances understanding of plant-microbiome interactions by providing baseline microbial data for developing beneficial bioengineering strategies and by assessing non-target impacts of engineered plant traits on the plant microbiome.

## Introduction

The biofuel industry has developed significantly over the past two decades given the impending need to replace fossil fuels, reduce greenhouse gas emissions, and mitigate climate change (Saladini et al., 2016; Alawan et al., 2019). Switchgrass (*Panicum virgatum* L.) is a C4 perennial grass, and is a flagship sustainable biofuel feedstock species in North America given its wide native range, fast-growth, high cellulose content, and relatively low requirements for water, nutrients and pesticides (Fike et al., 2006; Sanderson et al., 2006; Casler et al., 2015). Lignocellulosic material is the cheapest feedstock to produce biofuels (Huber et al., 2006), and nearly 80% of switchgrass dry weight biomass is composed of cellulose, hemicellulose, and lignin (David and Ragauskas, 2010).

Lignin is a major component of plant cell walls, and in grasses is composed of large branched and oxygenated polyaromatic compounds consisting of monomer units of coniferyl, sinapyl, and *p*-coumaryl alcohols (Evans et al., 1986; Huber et al., 2006). Since lignin contributes to biomass recalcitrance to deconstruction, reducing lignin content in feedstocks facilitates cellulose and hemicellulose hydrolysis, thus, increases fermentable sugar yields from biomass, and improves its conversion efficiency to bioenergy and advanced bioproducts (Sanderson et al., 2006; Xu et al., 2011; Ravindran and Jaiswal, 2016).

Several genetic engineering techniques have been used to reduce lignin content in plants (Liu and Eudes, 2022). These include the silencing of genes encoding lignin biosynthetic enzymes such as 4-coumarate: CoA ligase (Xu et al., 2011) and caffeate *O*-methyltransferase (Fu et al., 2011). Another promising strategy for reducing lignin in bioenergy crops involves the expression of bacterial 3-dehydroshikimate dehydratase (QsuB), which reduces the pool of precursors necessary for lignification (Eudes et al., 2015). In the shikimate pathway, 3-dehydroshikimate is the precursor of shikimate and phenylalanine, which are key metabolites involved in lignin biosynthesis (Tohge et al., 2013). QsuB converts 3-dehydroshikimate into protocatechuate and thereby impacts lignin synthesis. Such genetic modifications have been shown to improve the saccharification of biomass compared to wildtype plants (Eudes et al., 2015; Unda et al., 2022). For example, the expression of QsuB in switchgrass resulted in a 12-21% reduction in lignin content and 5-30% increase in saccharification efficiency, as well as greater bioaccumulation of protocatechuate (Hao et al., 2021).

Plant-associated microbiomes are composed of populations of diverse bacteria and fungi that colonize internal and external plant tissues, and may include beneficial, commensal, and pathogenic organisms. Microbiomes have been shown to be important to maintaining plant health and can be leveraged to increase biomass yield (Ghimire et al., 2009; Kim et al., 2012), enhance plant nutrient availability (Ker et al., 2014), improve drought tolerance (Ghimire and Craven, 2011), and further provide ecosystem services (Sher et al., 2020; Haan et al., 2023). For example, switchgrass plants inoculated with *Serendipita vermifera* (originally *Sebacina*) produced as much as 75% and 113% more shoot biomass at 2-month and 3.5-month harvest, respectively (Ghimire et al., 2009).

Plant genotype has been shown to be a factor involved in structuring plant microbiome, as was found for bacterial communities in the switchgrass rhizosphere, as well as aboveground and belowground fungal and bacterial microbiomes of switchgrass (Sutherland et al., 2022; Beschoren da Costa et al., 2022; Edwards et al., 2023). Similar results were found for switchgrass phyllosphere microbiomes in the field (VanWallendael et al., 2022). Changes in microbial community between highly productive and less productive switchgrass cultivars can be linked to the greater and lower microbial nitrogenase activity, respectively, which suggested a possible linkage between microbiomes and cultivar yields (Rodrigues et al., 2017; Ulbrich et al., 2021).

Genetic engineering can improve plant growth biomass and chemical properties, but it may also have unexpected impacts on plant-microbe interactions. For example, the silencing of cinnamoyl-CoA reductase gene reduced lignin in poplar trees, but also significantly changed the bacterial community in roots, stems, and leaves (Beckers et al., 2016). Similarly, poplar trees downregulated in genes encoding for the lignin biosynthetic enzymes caffeoyl-CoA O-methyltransferase, caffeic acid O-methyltransferase, cinnamoyl-CoA reductase, and cinnamyl alcohol dehydrogenase all displayed a lower mycorrhizal colonization in vitro (Behr et al., 2020). Thus, although growing engineered switchgrass with reduced lignin could have obvious industrial advantages regarding deconstruction and conversion processes, the impact of the engineered trait on the structure and functioning of the plant microbiome needs to be evaluated.

DeBruyn et al. (2017) reported that lower lignin lines of COMT (caffeic acid O-methyltransferase)-downregulated switchgrass plants had no effects on bacterial diversity, richness, or community composition of soil samples, but they did not investigate the fungal community and other plant compartments. In this study, we assessed in switchgrass the impact of the QsuB engineered plants on the microbiome across plant compartments. We accomplished this by characterizing both fungal and bacterial communities, within bulk soil, rhizosphere, root, leaf, and inflorescence of QsuB and wildtype Cave-in-Rock switchgrass. We hypothesized that QsuB engineered traits would alter the structure of fungal and bacterial microbiomes, particularly in belowground samples that support high amounts of microbial diversity.

## Method and Materials

### Plant Growth and Transplant

The transgenic switchgrass line *pZmCesa10*:*QsuB-5* and parental wildtype (cultivar Alamo-A4) used in this study have been described previously (Hao et al., 2021). Three transgenic and three wildtype plants were raised in axenic conditions and were then planted in sterile potting mix (Sure Mix, Michigan Grower Products Inc, Galesburg, MI, U.S.) and grown vegetatively (16-hr light and 8-hr dark at 23 ℃) to establish sufficient biomass to allow each plant to be split into three individuals. Deionized water was applied every other day and fertilizer was applied every other week. Splitting was done by excising each plant at the crown into three at approximately equal crown size with a sterilized scissor. The senesced aboveground tissues and old structural roots were trimmed off to only retain green aboveground tissue and minimum non-lignified young roots.

After splitting, switchgrass plants were planted into a blend of field soil and double-autoclaved play sand 50/50 (v/v). Field soil was collected from the top 20 cm of a switchgrass field in the long-term ecological research station for bioenergy cropping systems in Hickory Corner, MI in Aug. 2021, and was sieved through a 1-cm hardware cloth to homogenize and remove root fragments and organic debris before mixing with sterile sand. For microbiome analyses, nine biological replicates were used for both the wildtype and QsuB genotypes. Split plants were treated with the same plant care for three months before the final microbiome sampling and termination of the experiment.

### Sample Collection and Processing

Sampling of above and belowground switchgrass-associated microbiomes was carried out at two separate occasions: after splitting prior to planting in field soil (as pre-transplant sampling status), and three months after splitting and planting in the field soil (as post-transplant sampling status). Samples were collected from two soil niches (i.e. bulk, rhizosphere) and four plant niches (root endosphere, leaf, inflorescence, senesced leaves). Bulk soil from triplicate plant splits was collected with a sterile spatula avoiding root zones. Rhizosphere soil was sampled from each replicate by collecting three young lateral roots up to 3 cm with root hairs included from each plant. Roots were vigorously agitated to detach the loosely attached soil. The roots were then vortexed in ddH_2_O containing 0.05% Tween 20 for 20 min to dislodge the tightly attached soil. The washes were kept as rhizosphere soil samples, which contained both rhizosphere and rhizoplane communities. Washed roots were then surface sterilized in 6% hydrogen peroxide solution for 30 s and kept as root endophytic samples. Expanded young healthy leaves were sampled at splitting. Other aboveground tissues aside from young leaves including inflorescence and senesced leaves were also sampled at the end of the experiment by using sterile scissors. All aboveground tissues were sampled at 5 cm below the tips for approximately 1-cm from three randomly picked tissue objects.

### DNA Extraction and Illumina MiSeq Sequencing

Samples were flash frozen in liquid nitrogen within one hr after collecting. Samples were then freeze-dried with a SpeedVac (Thermo Fisher, Waltham, MA, U.S.), placed in 2-mL centrifuge tubes together with three metal beads in each tube, and ground to a powder with a TissueLyser II (Qiagen, Hilden, Germany) at maximum speed for 40 s. Microbial DNA was extracted from soil samples with a MagAttract PowerSoil DNA Kit (Qiagen, Hilden, Germany) and from plant samples with E.Z.N.A.® Plant DNA Kit (Omega Bio-Tek, Norcross, GA, U.S.). Libraries were prepared as previously described (Beschoren da Costa et al., 2022) with some modifications. Briefly, extracted DNA was amplified with primer sets 515f and 806r for bacterial communities targeting the 16S rDNA V4 region, and primers 5.8f and ITS4r for fungal communities targeting ITS2 rDNA (Caporaso et al., 2011; Taylor et al., 2016). Following the initial amplification, amplicons were PCR-ligated onto Illumina sequencing adapters and customized barcodes, and normalized with a Norgen DNA Purification Kit (Norgen Biotek Corp., Thorold, ON, Canada). Pooled barcoded amplicons were then purified and concentrated with Amicon centrifugal units (Sigma-Aldrich, St. Louis, MO, U.S.), and further purified with a HighPrep™ PCR Clean-up System (MAGBIO Genomics, Gaithersburg, MD, U.S.). Sequencing was conducted at Michigan State University RTSF Genomic Cores (East Lansing, MI, U.S.) with a v3 kit on a Illumina MiSeq sequencer. The raw sequences were demultiplexed with default setting in bcl2fastq, filtered, clustered into amplicon sequence variants (ASVs) using DADA2 (Callahan et al., 2016) in R 4.0.2. ASV taxonomic annotations were generated using CONSTAX2 v2.0.18 (Liber et al., 2021) with SILVA v138 (Quast et al., 2013) for the 16S and UNITE 9.0 (Abarenkov et al., 2010) for the ITS regions, respectively. Raw 16S and ITS sequences data were deposited in NCBI under BioProject ID PRJNA1002602 and PRJNA1002603, respectively.

### Statistical Analysis and Data Visualization

ITS and 16S rRNA amplicon sequence variant (ASV) tables, taxonomy tables, and metadata were imported into the R software for statistical computing and graphics. All ITS sequences with BLAST identity and coverage of ≤ 60% to the UNITE fungal database v 9.0 (Abarenkov et al., 2022) were removed from the dataset. Pre-transplant leaf samples and post-transplant senescence leaf samples were dominated by plant organelles (e.g. mitochondria, chloroplast) with a very low number of fungal and bacterial sequences, therefore, we removed these samples from our analysis (**Fig. S1** and **S2**). Mitochondria and chloroplast sequences were also removed from the overall 16S dataset. Sequence distributions allowed for the removal of outlier samples (one post-transplant non-QsuB leaf sample and one post-transplant non-QsuB root sample) with low fungal read counts were detected and deleted (**Fig. S3** and **S4**). Samples with low number of reads (i.e. distribution outliers) were removed by adopting rarefaction cutoffs of 2,948 and 12,126 sequence reads per sample for fungi and bacteria, respectively. Rarefaction curves were calculated in the *vegan* package (Oksanen et al., 2015) and plotted in the *ggplot2* package (Wickham, 2016). Rarefaction curves showed that most samples recover the whole diversity present in each sample, and rarefaction only marginally affected the total number of ASV detected across the entire datasets (**Fig. S5**).

Rarefied ASV richness and Shannon diversity index were calculated in *vegan*. Beta-diversity Bray-Curtis distance matrices were assessed to illustrate the community structures between samples and sample groups. We used non-metric multidimensional scaling (NMDS) and principal coordinate analysis (PCoA) ordinations to visualize beta-diversity. Permutational multivariate analysis of variance (PERMANOVA) was performed to test the statistical differences of beta-diversity between sample groups. Stacked bar charts were generated to show the relative abundance of lineage-level bacteria and fungi in sample groups. To identify differentially abundant ASV across sample groups we used a pairwise Wilcoxon test and DESeq2 in the *stats* (R Core Team, 2023) and *DESeq2* (Love et al., 2014) R packages. All analysis and figures were generated in R (R Core Team, 2023).

## Results

### Data Summary and Overview

In total, we obtained 30,086,564 16S and 17,811,594 ITS raw sequence reads, respectively, from the 198 sample libraries. After removing non-target and contaminants OTUs, a total of 24,577,163 16S and 13,403,151 ITS reads remained, respectively. These accounted for 33,853 and 8,089 OTUs for the 16S (bacterial) and ITS (fungal) communities, distributed across 144 total samples. On average, each sample had 338,995.4 (± 2,029,856 standard deviation) 16S sequence reads and 184,871 (± 1,106,116 standard deviation) ITS sequence reads.

The three experimental variables in our design were (1) status (pre-transplant and post-transplant to field soil); (2) niche (bulk soils, rhizosphere soils, roots, leaves, and inflorescences); and (3) and genotype (QsuB and non-QsuB wildtype). In the nonmetric multidimensional scaling (NMDS) analyses of fungal and bacterial datasets, samples from the same niche clustered together, especially in bacterial communities, when plotting on two dimensions (**Fig. S6**). Bulk soil and rhizosphere samples are visually close to each other in their fungal and bacterial communities. Bacterial communities of belowground samples (roots, rhizosphere and bulk soils) were distinct from those of aboveground samples (leaf and inflorescence) prominently. Pre- and post-transplant samples were also clearly separated in ordination space (**Fig. S6**).

Soils and sampling niches had obvious influences on both fungal and bacterial communities. Therefore, to investigate the influence of the genotype on microbial communities, we split our datasets by sampling niches and status in the following analysis.

### QsuB Leaf and Root Samples Had Higher Bacterial Richness and Alpha-diversity

In general, soil and rhizosphere samples had significantly higher richness than inflorescence and leaf samples. For fungal communities, root samples had relatively lower richness, compared to inflorescence and leaf samples. For bacterial communities, root samples had relatively higher richness than aboveground inflorescence and leaf samples, especially for post-transplant root samples (**Fig. 1**). The QsuB genotype had no significant influence on the fungal richness in any plant niches of pre-transplant or post-transplant samples (**Fig. 1A**, **1B**). However, QsuB plants had significantly greater bacterial richness in post-transplant leaf and pre-transplant root samples, but not in post-transplant root samples (**Fig. 1C**, **1D**).

**Figure 1.**
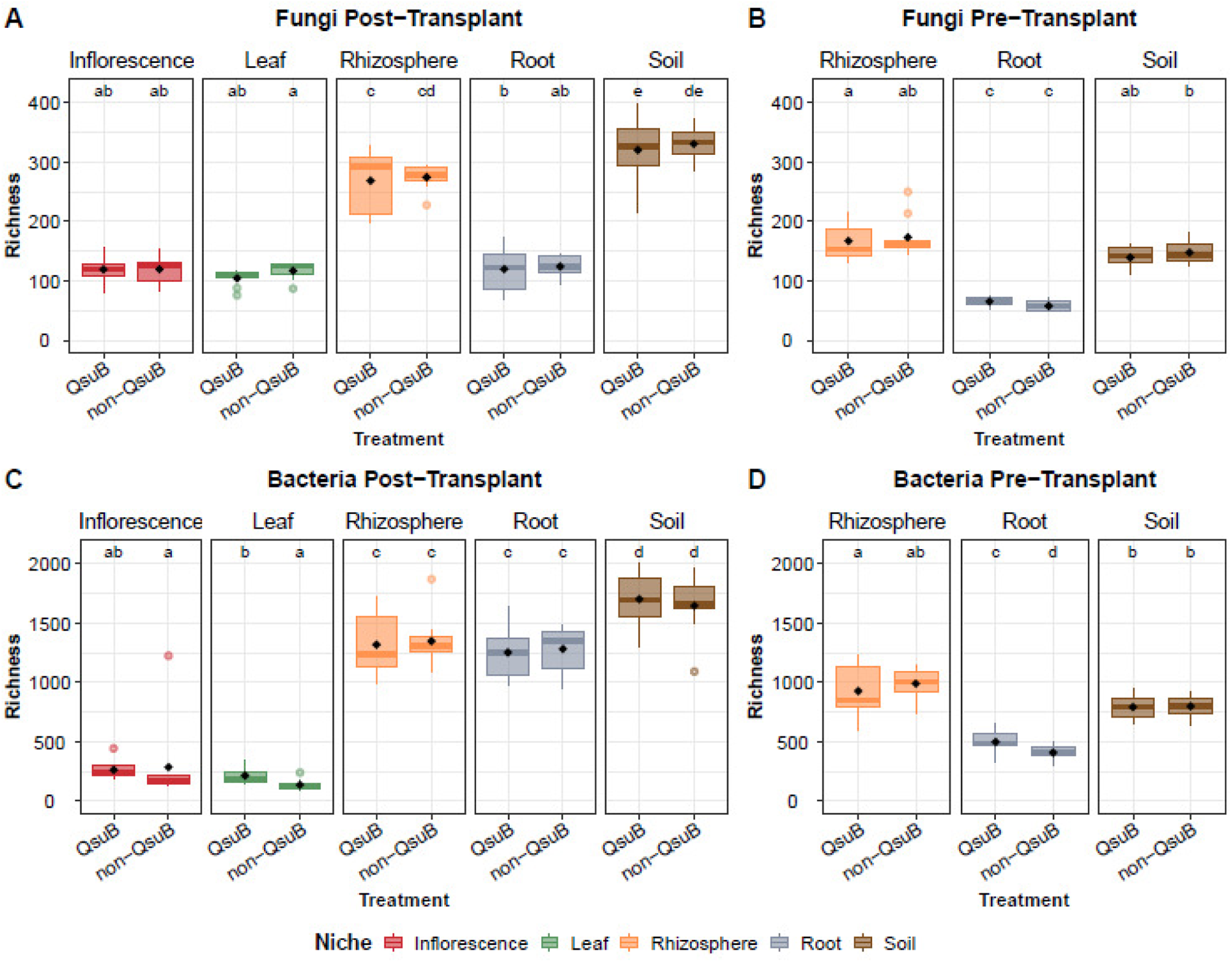
Boxplot of observed ASV richness for Post-Transplant Fungi (A), Pre-Transplant Fungi (B), Post-Transplant Bacteria (C), and Pre-Transplant Bacteria (D) grouped by sampling niches (inflorescence, leaf, rhizosphere, root, and soil). Letters represent pairwise Wilcoxon tests among groups (*p ≤ 0.05* after Bonferroni adjustment).

Fungal communities in root samples had significantly lower Shannon diversity compared to soil, rhizosphere and aboveground tissues. The QsuB genotype had no significant influence on the fungal Shannon diversity indices across sampling niches for both pre-transplant and post-transplant samples (**Fig. 2A**, **2B**). For bacterial communities, soil samples (soil and rhizosphere) had significantly greater diversity than plant samples (root, inflorescence, and leaf) (**Fig. 2**). The QsuB plants had significantly greater bacterial Shannon indices in post-transplant inflorescence, leaf, and pre-transplant root samples, but not in post-transplant root samples (**Fig. 2C**, **2D**).

**Figure 2.**
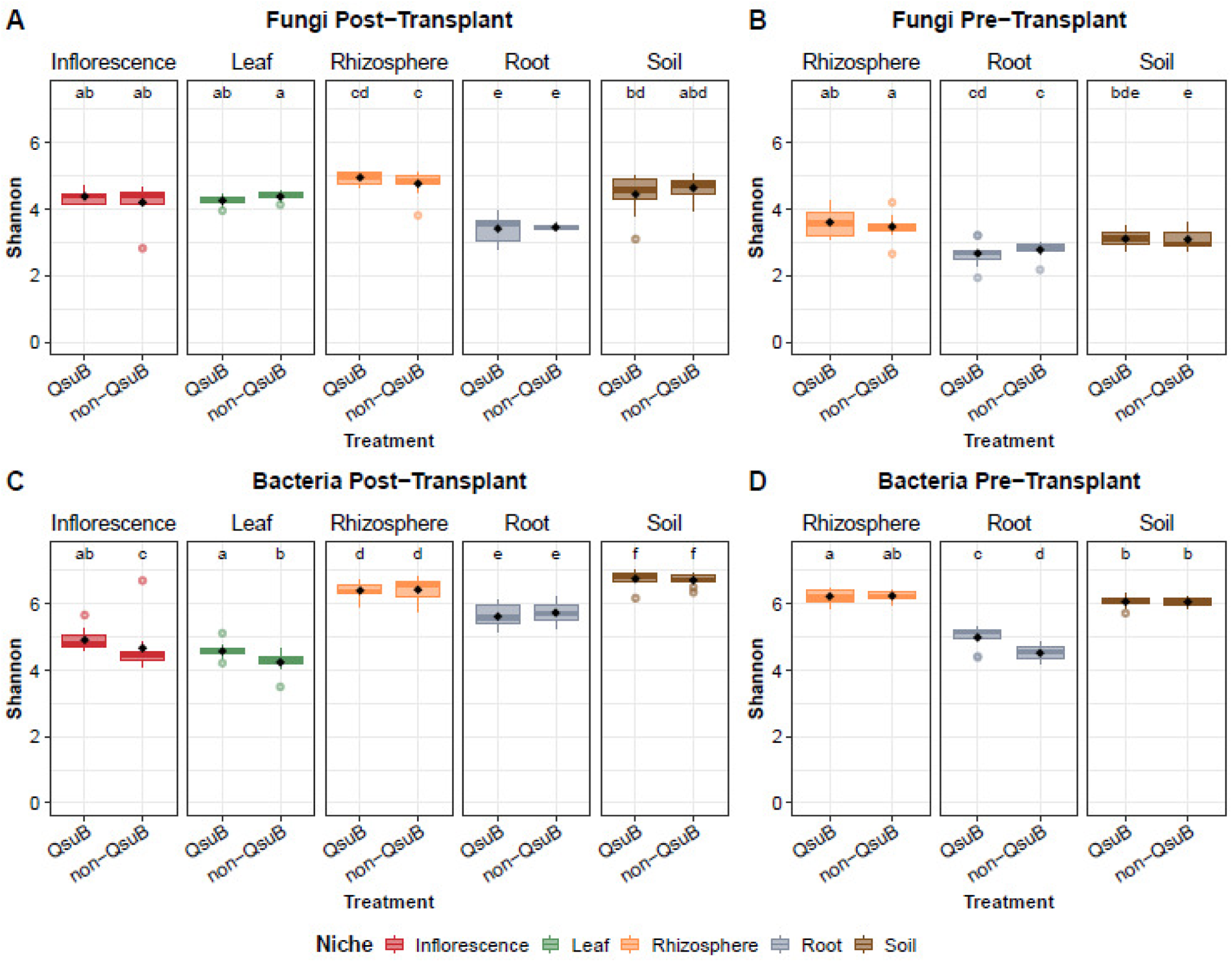
Boxplot of Shannon indices for Post-Transplant Fungi (A), Pre-Transplant Fungi (B), Post-Transplant Bacteria (C), and Pre-Transplant Bacteria (D) grouped by sampling niches (inflorescence, leaf, rhizosphere, root, and soil). Letters represent pairwise Wilcoxon tests among groups (*p ≤ 0.05* after Bonferroni adjustment).

Additionally, it is worth noting that, for the same sampling niches, post-transplant samples always had greater bacterial and fungal richness and Shannon indices than those of corresponding pre-transplant samples (**Fig. 1** and **2**).

### QsuB Significantly Influenced Root and Leaf Fungal Community Beta-diversity

We used principal coordinate analysis (PCoA) ordinations to improve visualization of beta-diversity results (**Fig. 3A**) and statistically examined the treatment effects on the beta-diversity. The QsuB traits significantly influenced the fungal community structures in the root (*p = 0.002*) and post-transplant leaf (*p = 0.041*) samples according to PERMANOVA (**Table S1**). In root samples, genotype, status, and the interaction between them were all significant factors of the fungal community structures. The QsuB genotype explained the most variance with the highest R^2^ of 23.09% (*p = 0.002*), followed by status (R^2^ = 17.68%, *p = 0.002*) and the interaction (R^2^ = 6.47%, *p = 0.003*), with residual of 52.76% (**Table S1**).

**Figure 3.**
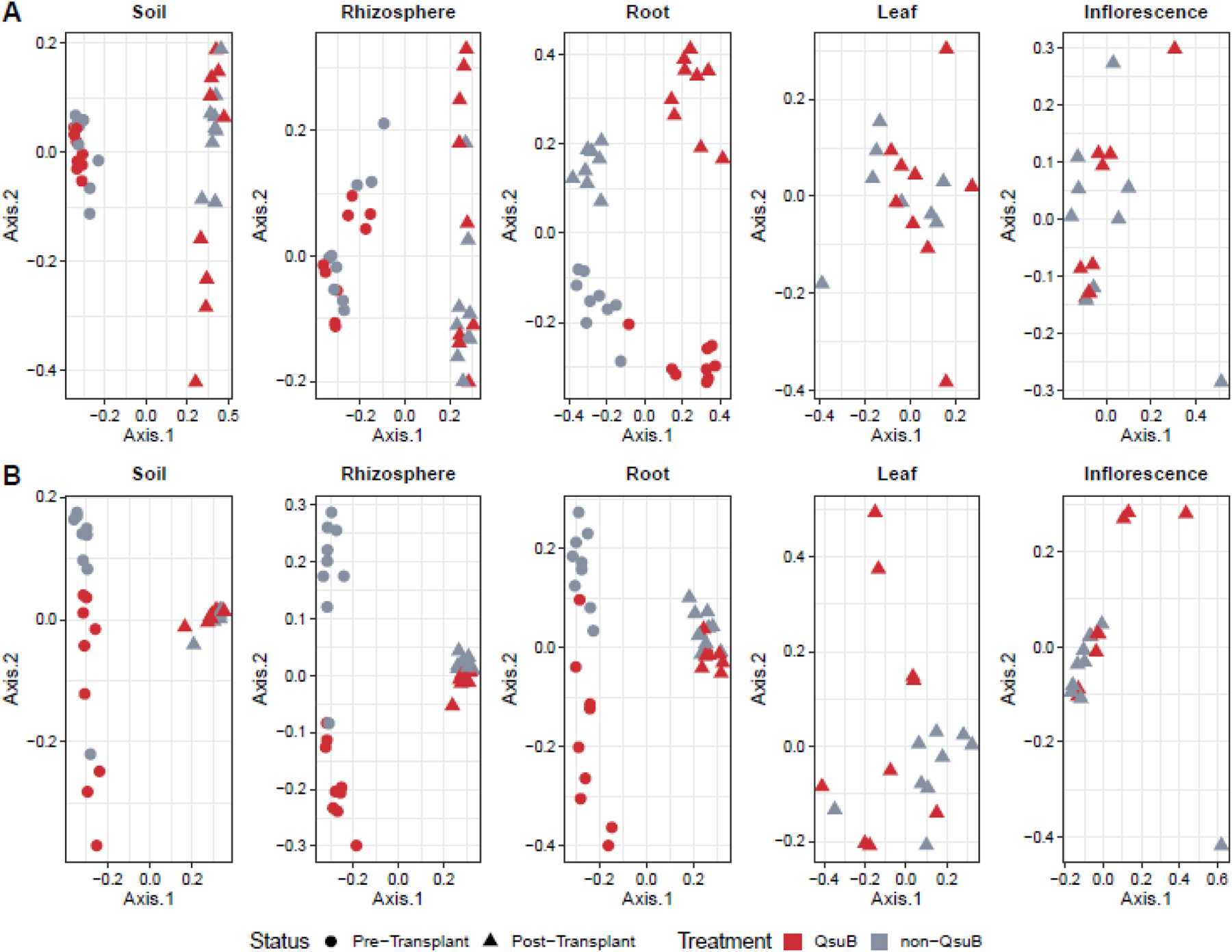
Principal coordinate analysis (PCoA) ordinations of fungal (A) and bacterial (B) communities sampled from soil, rhizosphere, root, leaf, and inflorescence niches at Pre-Transplant and Post-Transplant.

### QsuB Significantly Influenced Root and Rhizosphere Bacterial Community Beta-diversity

The root bacterial community of QsuB genotype and wildtype switchgrass separated from each other on two-dimension PCoA (**Fig. 3B**) and the visual observation was supported by the PERMOVA. QsuB genotype significantly influenced the bacterial community structures in root (*p = 0.003*) and rhizosphere (*p = 0.029*) samples (**Table S1**). In both root and rhizosphere samples, genotype, status, and the interaction between them were all significant factors of the bacterial community structures. In root samples, the status explained the most variance with R^2^ of 20.56% (*p = 0.003*), followed by QsuB genotype (R^2^ = 5.67%, *p = 0.003*) and their interaction (R^2^ = 4.35%, *p = 0.019*), with residual of 69.42% (**Table S1**). In rhizosphere samples, the status also explained the most variance with R^2^ of 21.31% (*p = 0.003*), followed by QsuB genotype (R^2^ = 4.15%, *p = 0.029*) and their interaction (R^2^ = 3.83%, *p = 0.029*), with residual of 70.71% (**Table S1**).

One thing to note is that, for both root and rhizosphere samples, the influence from the QsuB genotype was only manifested in pre-transplant samples in the PCoA plots, but not in post-transplant samples, which seemed clustered together (**Fig. 3B**). To focus on the impact of the QsuB genotype and eliminate interference from different status, we ran beta-diversity analysis only on the post-transplant root and rhizosphere samples. This approach revealed that the QsuB genotype had significant influence on the bacterial community of post-transplant root (*p = 0.001*) and rhizosphere (*p = 0.003*) samples (**Fig. S7**, **Table S1**).

### Fungal Composition

Overall, sixteen fungal lineages were detected in our sample, with *Ascomycota*, *Glomeromycotina*, and *Basidiomycota* being predominant and averaging relative abundance of 80.63%, 8.57%, and 7.97%, respectively (**Fig. S8**). Most *Glomeromycotina* were detected in roots samples and the majority of *Basidiomycota* were detected in post-transplant bulk soil and rhizosphere samples (**Fig. S8**). The above beta-diversity analyses showed that the QsuB genotype significantly influenced the fungal community of root and leaf samples. Around 94.00% of leaf fungal are *Ascomycota*, providing little information at the lineage-level fungal composition (**Fig. S8**). At order-level, QsuB plants had relatively more *Hypocreales* compared to the wildtype in leaves (**Fig. 4**), but the difference was not statistically significant.

**Figure 4.**
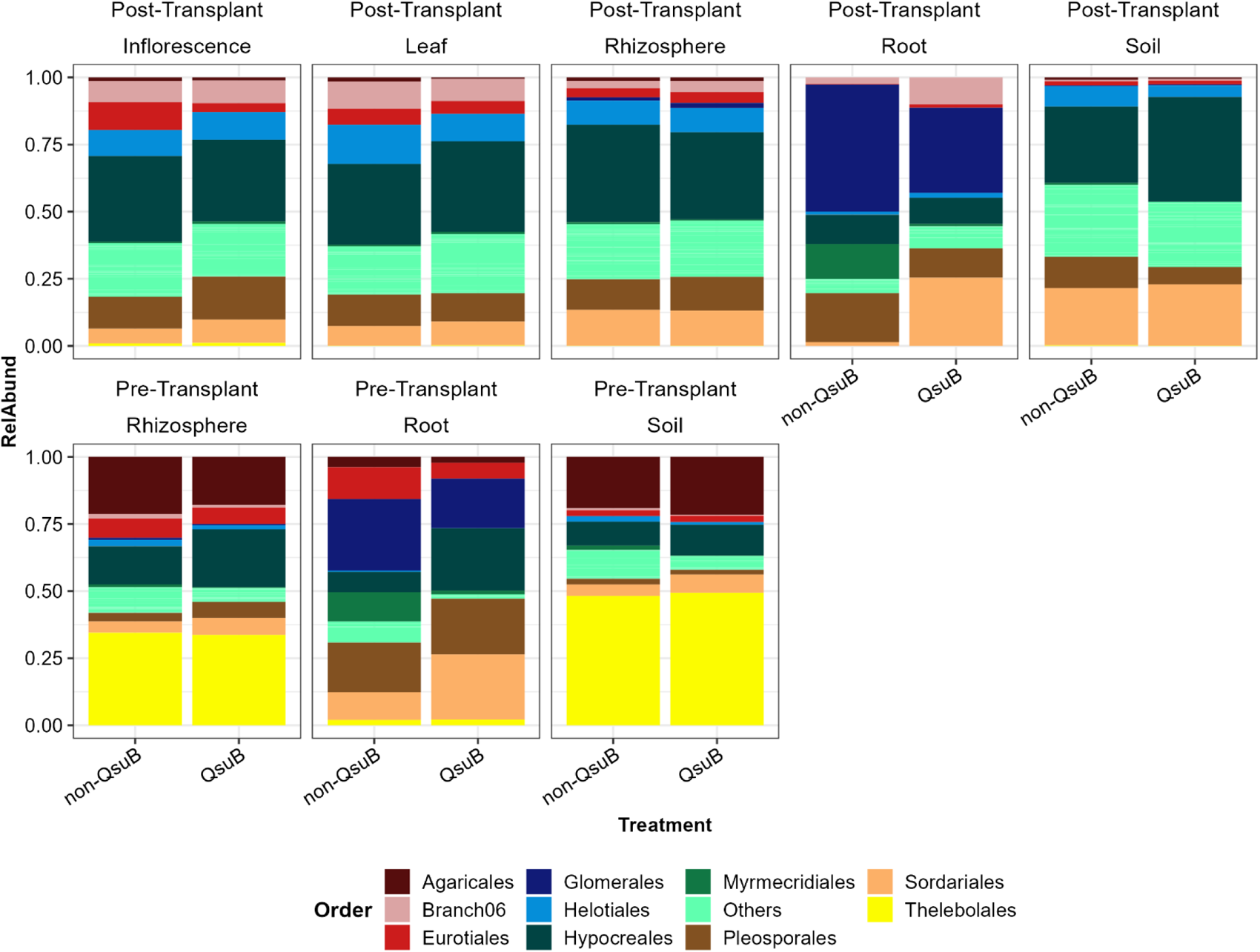
Order-level fungal taxonomic distribution of samples from inflorescence, leaf, rhizosphere, root, and bulk soil niches of Post-Transplant and Pre-Transplant.

The significant effects of QsuB genotype on root fungal communities were likely explained by relatively more *Ascomycota* and relatively fewer *Glomeromycotina* in QsuB plants (**Fig. S8**). In post-transplant root samples, QsuB plants had 62.52% *Ascomycota* and 32.86% *Glomeromycotina*, while non-QsuB wildtype had 50.74% *Ascomycota* and 47.76% *Glomeromycotina*. Statistical analysis on lineage-level relative abundance data showed that only *Ascomycota* (*p < 0.001*) were significantly influenced by QsuB genotype in post-transplant root samples. However, in pre-transplant root samples, both *Ascomycota* (*p = 0.009*) and *Glomeromycotina* (*p = 0.033*) were significantly influenced by QsuB genotype. In pre-transplant root samples, QsuB plants had 78.35% *Ascomycota* and 18.57% *Glomeromycotina*, while non-QsuB wildtype had 67.63% *Ascomycota* and 27.71% *Glomeromycotina* (**Fig. S8**).

To further investigate the QsuB genotype effects, we performed the relative abundance of fungi at order level. We listed the top 10 (by abundance) orders and summed up other orders as “Others” (**Fig. 4**). In pre-transplant root samples, 3 of top 10 orders were significantly influenced by QsuB genotype and they are all from class *Sordariomycetes*. QsuB plants had significantly more *Hypocreales* (*p = 0.021*) and *Sordariales* (*p = 0.010*), but less *Myrmecridiales* (*p = 0.005*). Obviously, much more *Sordariales* and less *Myrmecridiales* in QsuB plants were also observed in post-transplant root samples. In pre-transplant samples, the relative abundance of *Glomerales* decreased from 27% in non-QsuB to 18% in QsuB, while in post-transplant samples, it decreased from 47% to 32% (**Fig. 4**).

### Bacterial Composition

Overall, 47 bacterial lineages were detected in our samples and the sum of top 10 most abundant lineages accounted for an average relative abundance of 95.64% among all samples. *Proteobacteria* and *Actinobacteria* were predominant bacterial lineages in our samples, with the average relative abundance of 45.47% and 17.53%, respectively (**Fig. 5**).

**Figure 5.**
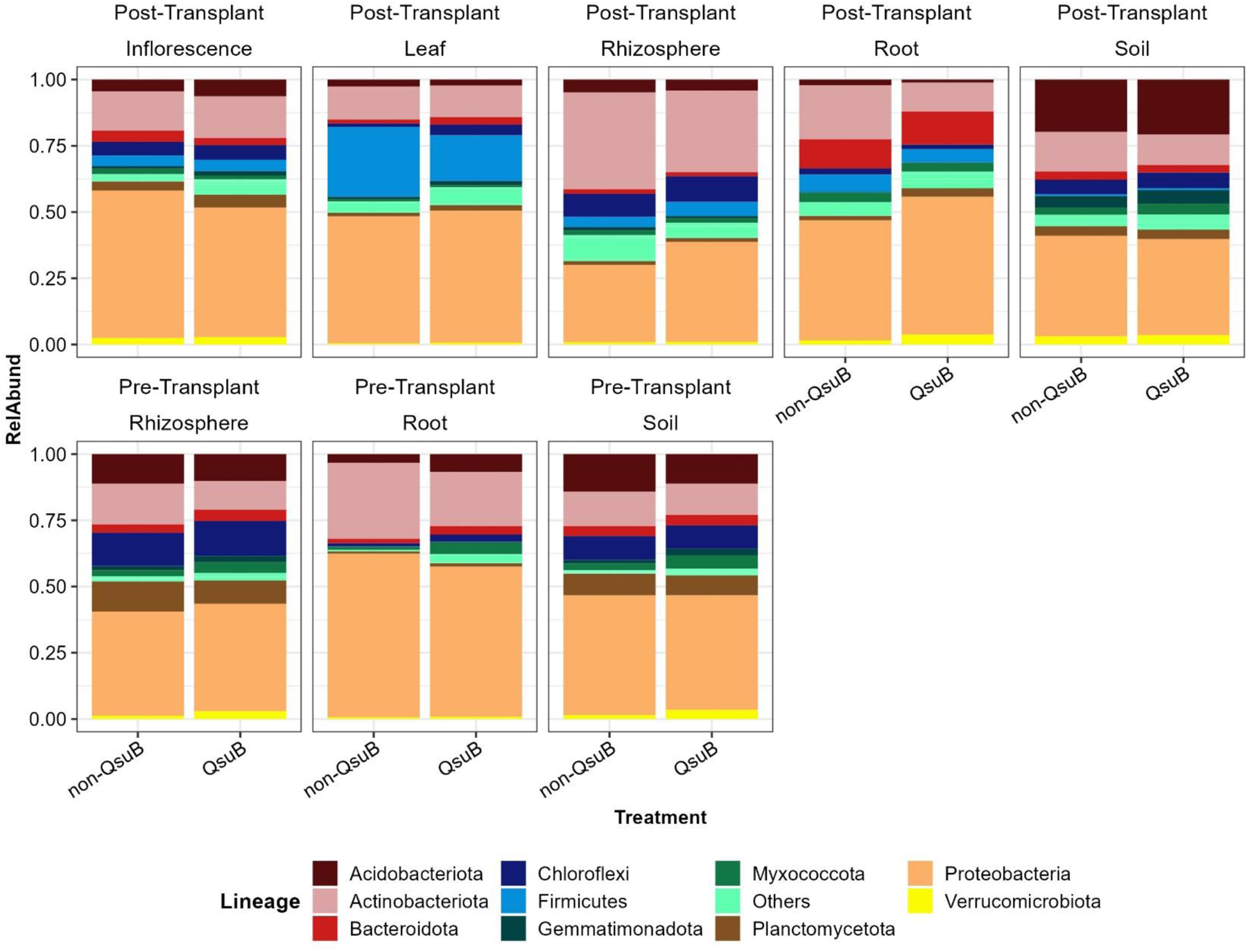
Lineage-level bacterial taxonomic distribution of samples from inflorescence, leaf, rhizosphere, root, and bulk soil niches of Post-Transplant and Pre-Transplant.

**Figure 6.**
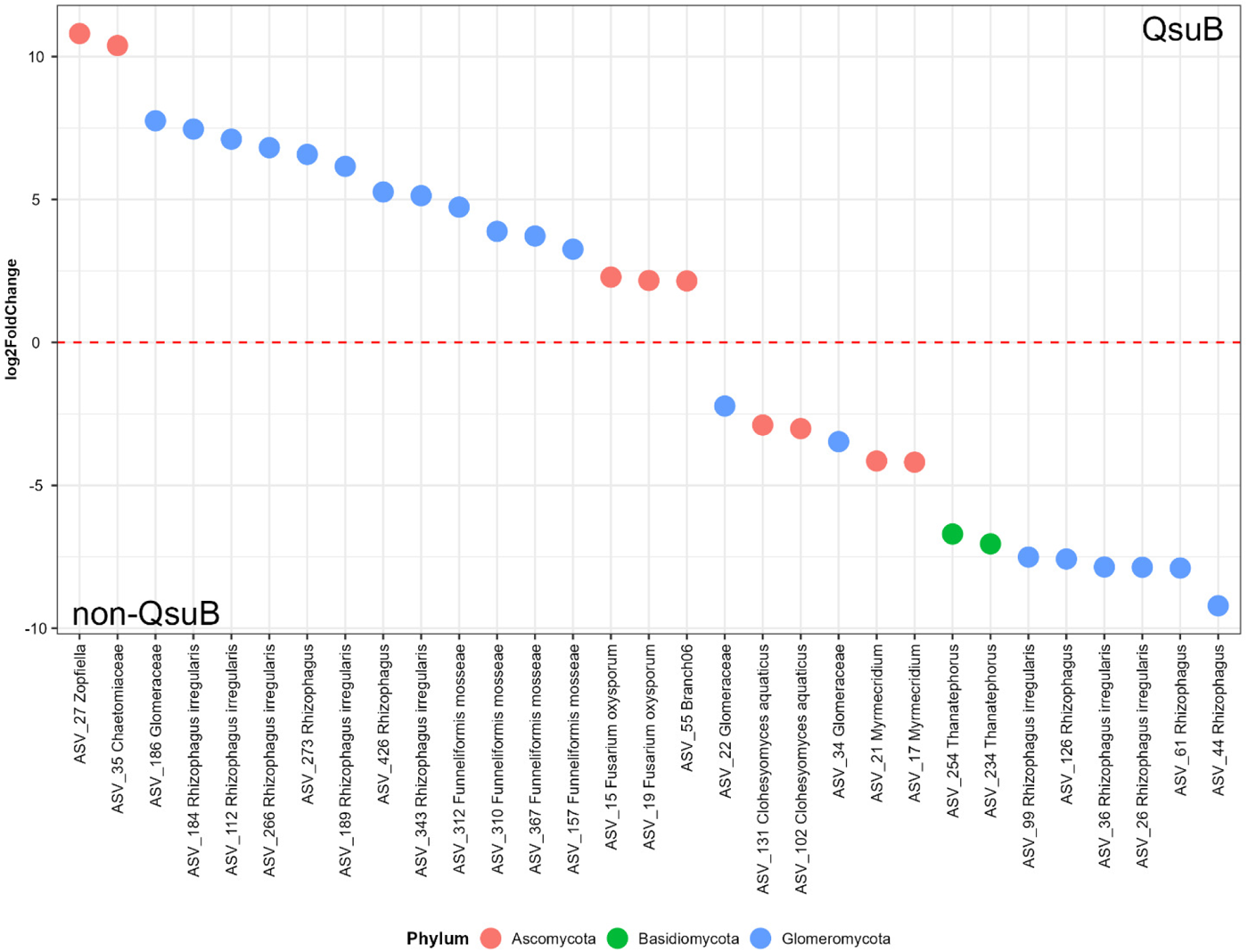
Differential expression analysis of fungal Post-Transplant root samples by DESeq2 displays the fungal amplicon sequence variants (ASVs) that are significantly differentially abundant between QsuB and non-QsuB wildtype plants. Different colors represent different phyla each sample belongs to. The ASVs above the dash line (log_2_FoldChange > 0) are significantly more abundant in the QsuB traits, while the ASVs below the dash line (log_2_FoldChange < 0) are significantly more abundant in the non-QsuB wildtype.

Bacterial communities of root and rhizosphere were significantly influenced by QsuB genotype. The only consistent trend associated with bacterial community was QsuB plants having relatively fewer *Actinobacteria* than wildtype: 20.38% vs. 28.72% (pre-transplant root), 10.93% vs. 20.45% (post-transplant root), 10.90% vs. 15.41% (pre-transplant rhizosphere), and 30.80% vs. 36.56% (post-transplant rhizosphere) (**Fig. 5**). In pre-transplant rhizosphere and root samples, we observed relatively more *Myxococcota* in QsuB than wildtype, but this trend was not shown in post-transplant samples; while in post-transplant rhizosphere and root samples, we observed relative more *Proteobacteria* in in QsuB than wildtype, but this trend was not shown in pre-transplant samples (**Fig. 5**). All of these trends were not statistically significant.

The bacterial relative abundance of the top 10 orders is shown in **Fig. S9**. Some trends are numerically evident: compared to wildtype, QsuB plants had relatively more *Pseudonocardiales* in roots and more *Burkholderiales* in rhizosphere (**Fig. S9**). The greater alpha-diversity (richness and Shannon index) in QsuB plants than wildtype was likely explained by relatively more “Others” in pre-transplant root samples (28.90% in QsuB vs. 14.02% in wildtype), in post-transplant inflorescence samples (45.27% in QsuB vs. 37.30% in wildtype), and in post-transplant leaf sample (30.21% in QsuB vs. 27.71% in wildtype) (**Fig. S9**).

### Indicator ASVs

We used differential abundance measurements between QsuB genotype and wildtype switchgrass to identify indicator ASVs. Consistent with our results, the Wilcoxon test commonly outputs a high number of significant ASVs compared to DESeq2 since the former is less stringent (Nearing et al., 2022; Botos et al., 2023). DESeq2 identified 14 and 15 bacterial biomarkers in pre- and post-transplant root samples, respectively, while Wilcoxon test identified 99 and 307 significant ASVs in those metadata subgroups. We barely found indicator ASVs in leaf fungal and rhizosphere bacterial samples with DESeq2, even though those samples’ microbial community were significantly influenced by QsuB genotype (**Fig. 3**). Root samples were the only group (by niche) that were significantly influenced in both fungal and bacterial communities (**Fig. 3**), so we focused on the fungal and bacterial indicator ASVs of root samples. Most indicator ASVs identified with DESeq2 were also identified with the Wilcoxon test.

Thirty-one and thirty-two fungal indicator ASVs were identified in the pre- and post-transplant root samples, respectively, the majority of which (14 of 31 from pre-transplant and 20 of 32 from post-transplant) were *Glomeromycotina* (**Fig. S10** and **6**). All of the identified *Glomeromycotina* belonged to the arbuscular mycorrhizal fungi (AMF) family *Glomeraceae*. Four *Funneliformis* (AMF) indicator ASVs were identified in post-transplant roots but not in pre-transplant root samples.

Fifteen indicator ASVs were detected both in pre- and post-transplant root samples. Six of them were associated with the QsuB genotype: ASV_27 (*Zopfiella*), ASV_35 (*Chaetomiaceae*), and ASV_112, 184, 189, 273 (*Rhizophagus*). Nine of them were associated with the wildtype switchgrass: ASV_21 (*Myrmecridium*), ASV_22, 34 (*Glomeraceae*), and ASV_26, 36, 44, 61, 99, 126 (*Rhizophagus*) (**Fig. S10** and **6**).

Compared to the fungal indicator ASVs, much fewer bacterial indicator ASVs were identified by DESeq2. Bacterial indicator ASVs were dominated by *Proteobacteria* and *Actinobacteriota* in pre- and post-transplant root samples, respectively. In pre-transplant root samples, *Lentzea aerocolonigenes* (ASV_191 and 244) were associated with the QsuB plants, while *Amycolatopsis mediterranei* (ASV_46 and 69) were associated with non-QsuB wildtype plants (**Fig. S11**). Both genera belong to *Pseudonocardiales*. In post-transplant root samples, *Streptomyces* (ASV_101) was the only *Actinobacteria* indicator associated with the QsuB plants (**Fig. S12**).

## Discussion

In this research we set out to assess whether switchgrass engineered for low lignin with the QsuB gene would impact the microbiome associated with different aboveground and belowground plant organs. As we hypothesized, our results indicate that QsuB engineered plants impacted switchgrass-associated microbial community structure. Our results showed that the QsuB genotype influenced fungal and bacterial community structure. Specifically, QsuB plants had a significant impact on the fungal community in root and leaf samples, and also a significant impact on belowground bacterial microbiomes in the root and rhizosphere. In contrast, we observed little impact of the QsuB genotype on inflorescence and bulk soil fungal or bacterial microbiomes.

### QsuB Plants Showed Lower Relative Abundance and Diversity of AMF

Arbuscular mycorrhizal fungi (AMF) are obligate biotrophic plant mutualists the belong to *Glomeromycotina* and are known to be beneficial to plant nutrition and soil health by transporting nutrients (e.g. P and N) and water to plant hosts via their hyphal network, while stimulating and stabilizing soil organic matter (Mosse et al., 1973; Leigh et al., 2009; Hestrin et al., 2022; Li et al., 2023; Nottingham et al., 2013). Under nutrient limitation, host plants are more dependent on AMF for nutrients (Kleikamp and Joergensen, 2006; Smith and Smith, 2011). In this study we detected fewer *Glomerales* (i.e. the most frequent AMF order detected in this study) sequences in the root samples from QsuB plants (both pre- and post-transplant).

The expression of QsuB in switchgrass results in the reduction of lignin and the accumulation of protocatechuate in biomass, and improves biomass saccharification efficiency (Hao et al., 2021). Less lignin and more protocatechuate may have stimulated bacterial community activity, and increased soil nutrient mineralization and turnover rates (De Graaff et al., 2010), which in turn may affect AMF colonization. Five lower lignin transgenic lines of poplar with downregulated for genes of monolignol biosynthesis pathway displayed a lower mycorrhizal colonization percentage than wildtype, and the authors proposed that the gene modifications in monolignol pathway impacted ectomycorrhizal colonization possibly by changing cell wall ultrastructure and decreasing the communication efficiency between plants and fungi (Behr et al., 2020).

### QsuB Roots Showed an Increase in the Relative Abundance of *Sordariales* and *Hypocreales*

In this study, more *Ascomycota* (e.g. *Sordariales* and *Hypocreales*) were detected in the root samples from QsuB plants compared to the wildtype. Previous work has identified *Sordariales* and *Hypocreales* as dominant decomposers in arable soil with long-term organic management practice (Ma et al., 2013), and we found these orders predominant in our samples. It is known that mycorrhizal and saprotrophic fungi compete for niche space and organic substrates (Boddy, 2000; Leifheit et al., 2015; Bödeker et al., 2016). For example, Cao et al. (2022) reported that AMF inhibited the population abundance and enzyme activity of saprotrophic fungi, possibly by reducing the availability of limiting nutrients. Increased accessibility to carbohydrates and soil nutrients may have stimulated *Sordariales* and *Hypocreales*, which were associated with the QsuB genotype.

Interestingly, in both root and leaf samples, QsuB plants had a higher relative abundance of *Hypocreales*. Six *Fusarium* (members of Hypocreales) ASVs were identified as indicator ASVs. The relative abundances of these *Fusarium* indicator ASVs significantly increased in the root and leaf samples of QsuB plants compared to the wildtype. This is of interest because many *Fusarium* species are known plant pathogens (Collins, 2018). In root samples, QsuB plants had relatively more *Fusarium* than the wildtype: 9.81% vs. 2.82% (pre-transplant) and 7.27% vs. 1.97% (post-transplant); however, this trend was not obvious in the leaf samples (**Fig. S13**). The only two leaf *Fusarium* indicator ASVs were identified at the species level: *Fusarium oxysporum* (ASV_15 and 19), and they were also root *Fusarium* indicator ASVs. Soils are often the source of plant-associated *F. oxysporum* (Dhaya, 2020), and detached leaf assays showed that *F. oxysporum* might be benign or beneficial in switchgrass, even though other *Fusarium* species were pathogenic (Collins, 2016). In this study, *F. oxysporum* was the only *Fusarium* species identified in leaf samples associated with QsuB plants. Knowing the diversity of *Fusarium* species, the complexity of their function, and the limits of short amplicon sequencing, the actual roles of the identified *Fusarium spp.* in our study are difficult to discern (Chandra et al., 2011).

In root samples, there were significantly more *Sordariales* in switchgrass QsuB plants than the wildtype, and this might be mainly contributed by *Zopfiella* (**Fig. S13**). *Zopfiella* spp. has the potential to control plant disease by producing antifungal compounds and promote plant growth by increasing stress resistance (Huang et al., 2015; Zhong et al., 2022).

### QsuB Plants Hosted a Greater Richness and Diversity of Bacteria

We found that QsuB engineered plants supported a significantly greater richness and diversity of bacteria in inflorescence, leaf, and root (pre-transplant) samples. Generally, bacteria are efficient at degrading simple substrates, while fungi are better equipped at decomposing recalcitrant organic matter, such as lignin (Boer et al., 2005). Lignin biodegradation starts with lignin depolymerization, which is predominantly performed by fungi (Kamimura et al., 2019).

Therefore, compared to lignin, protocatechuate represents a more favorable growth substrate for bacteria to utilize, and bacteria are competitive for carbon and energy sources. Plants expressing QsuB accumulate inside tissues more protocatechuate that may stimulate the activity of the bacterial community. Diverse bacteria are indeed known to degrade protocatechuate including those in the order *Bacillales*, *Burkholderiales*, *Sphingomonadales*, and *Pseudomonadales* (Noda et al., 1990; Hardwood and Parales, 1996; Kamimura et al., 2010; Kamimura and Masai, 2014).

### Fewer *Actinobacteria* Were Detected in the Root and Rhizosphere Samples of QsuB Plants

In this study, the relative abundance of *Actinobacteria* detected in the root and rhizosphere samples of QsuB plants was lower than that of the wildtype. *Actinobacteria* are an important terrestrial group of detritus decomposers (Boer et al., 2005), which are able to degrade lignin materials (Trigo and Ball, 1994). Given that QsuB engineered switchgrass biosynthesizes less lignin, it may be expected to host a lower relative abundance of *Actinobacteria*. In post-transplant root samples, all non-QsuB associated bacterial indicator ASVs showing significantly increased relative abundances were *Actinobacteria* (**Fig. S9**).

### The AMF and Bacterial Community Dynamics May Be Interlinked

We have described how the QsuB genotype influenced the fungal communities, especially AMF, as well as the bacteria communities belowground. We posit that the change in the AMF community might be an important driver of the change observed in those bacterial communities. Previous studies have found that the AMF community composition was a significant contributor to determining the bacterial community composition (Rillig et al., 2006), perhaps through changes in the root exudates composition and soil structure modification (Barea et al., 2005; Boer et al., 2005). Interestingly, AMF-associated bacterial communities have been shown to be structured predominantly by AMF symbiont identity (*Glomus geosporum* or *Glomus constrictum*), rather than the host plant (*Plantago lanceolata* or *Hieracim pilosella*) (Roesti et al., 2005). AMF hyphae release a variety of exudates, including carbohydrates, polyols, amino acids, amines, nucleic acids, organic acids, etc.; different AMF species or the same AMF under different abiotic conditions might have different metabolite profiles of hyphal exudates (Luthfiana et al., 2021; Faghihinia et al., 2023). The carbon sources supplied by AMF have important roles in bacterial growth and distribution, so it is likely that AMF activity has an impact on the surrounding bacterial communities (Zhou et al., 2020; Jiang et al, 2021; Faghihinia et al., 2023). AMF hyphae provide a scaffold bridging the soil and root microbiomes (Wang et al., 2023). The addition of protocatechuate to the culture medium inhibited primary root growth but increased lateral root numbers in Arabidopsis (Herbert et al., 2019). The potential root morphology change may also influence AMF development and surrounding bacterial communities.

Some specific bacteria showed the same (positive or negative) response to AMF, even under different experimental setups. For example, in this study, root and rhizosphere samples from QsuB plants had relatively fewer AMF and *Actinobacteria*. This positive response of *Actinobacteria* to AMF has also been observed in other studies (Frey-Klett et al., 2007; Faghihinia et al., 2023). Recent work showed that QsuB switchgrass had no yield penalty compared to the wildtype under optimal irrigation in the field (Eudes et al., 2023). However, as AMF and *Actinobacteria* are associated with plant drought resilience, this raises questions into how QsuB plants would fare under water-limiting conditions (Xu et al., 2018; Chandrasekaran, 2022).

### Importance of Testing Microbiome Impacts in QsuB Engineered Plants

Engineering the biofuel feedstock switchgrass with the QsuB gene is a promising strategy for reducing lignin content and improving saccharification. Our work highlights the importance of assessing QsuB genotype impact on the plant-associated microbiomes. We found that QsuB engineering changed plant physiology and its microbiomes, including some important functional microbial groups including AMF and *Actinobacteria*. Unfortunately, we did not obtain chemical data of the roots and surrounding rhizosphere, so the lignin and protocatechuate contents of specific belowground compartments is unknown. It is possible that the accumulation of protocatechuate altered plant cell osmolarity, and might further modify the root exudates, which may directly contribute more to the microbial community dynamics. A longer-term field study of QsuB bioenergy plants with measurements of biomass yield, soil properties, and microbiome communities in different locations and climates could help confirm the promise and future of QsuB-engineered bioenergy crops under real-world agricultural scenarios.

## Conclusion

Less lignin content, more fermentable sugar yield, and greater biomass conversion efficiency to biofuels made QsuB genotype promising in wide application. As hypothesized, our study found that the QsuB engineered plants impacted switchgrass associated fungal and bacterial communities, especially those from roots and the rhizosphere. Importantly, the microbiome differences between QsuB plants and non-modified wildtype switchgrass did not appear to impact the abundances of putative switchgrass pathogens. However, the reduction in AMF diversity and abundance in QsuB plants are noteworthy and raise questions regarding how this could further impact plant performance under drought conditions, and consequent soil physio-chemical properties. By characterizing the microbiome responses to QsuB genotype we provide a baseline for evaluating QsuB and other bioengineered traits on plant-microbe interactions.

## Acknowledgements

We thank David Lowry and Linnea Fraser for helpful discussions and assistance in switchgrass splitting and plant care. We acknowledge Michigan State University RTSF Genomics Core for generating sequencing data from our amplicon libraries. This material is based upon work supported by the Great Lakes Bioenergy Research Center, U.S. Department of Energy, Office of Science, Biological and Environmental Research Program under Award Number DE-SC0018409. Part of this material is based upon work supported by the Joint BioEnergy Institute, U.S. Department of Energy, Office of Science, Biological and Environmental Research Program under Award Number DE-AC02-05CH11231 with Lawrence Berkeley National Laboratory.

